# Mutations differentially affecting the coronavirus Mac1 ADP-ribose binding and hydrolysis activities indicate that it promotes multiple stages of the viral replication cycle

**DOI:** 10.1101/2025.04.08.647734

**Authors:** Joseph J. O’Connor, Anuradha Roy, Reem Khattabi, Catherine Kerr, Nancy Schwarting, Yousef M. Alhammad, Philip Gao, Xiaoming Zhang, Xufang Deng, Anthony R. Fehr

## Abstract

All coronaviruses (CoVs) encode a conserved macrodomain, termed Mac1, in non-structural protein 3 (nsp3) that binds and hydrolyzes ADP-ribose covalently attached to proteins. Mac1 is a key virulence factor that counters antiviral ADP-ribosyltransferase (PARP) activity. Previously, we found that MHV strain JHM (JHMV) with a mutation in the adenine binding site, JHMV-D1329A, was extremely attenuated in all tested cell types as opposed to JHMV-N1347A, which only has a replication defect in bone-marrow derived macrophages (BMDMs). Interestingly, an N1347A/D1329A double mutant was unrecoverable, indicating an essential role for Mac1 in JHMV infection. We hypothesized that these mutations may impact different stages of the MHV lifecycle. First, to clarify how these mutations affected the biochemical activities of Mac1, we generated Mac1 proteins encoding the same mutations. As expected, the D-A mutation was extremely defective in ADP-ribose binding but maintained enzyme activity. In contrast, we previously found that the N-A mutation had WT levels of ADP-ribose binding but low enzyme activity, confirming that these mutations differentially effect the biochemical functions of Mac1. Following infection, D1329A displayed a large defect in the accumulation of viral RNA compared to WT or N1347A in all cells tested. Alternatively, N1347A infection produced normal levels of viral RNA but produced reduced levels of viral protein in interferon competent bone-marrow derived macrophages (BMDMs). These results suggest that Mac1 ADP-ribose binding and enzymatic activities promote different stages of the viral lifecycle, demonstrating the critical importance of Mac1 for JHMV replication.

**IMPORTANCE:** Over the last three decades, coronaviruses have repeatedly demonstrated their potential to become significant veterinary and public health threats. Zoonotic transmission of the myriad known coronavirus strains will remain a concern, regardless of the advances in vaccines and treatment. One difficulty in anticipating the next coronavirus outbreak is their diverse lineage and high propensity for mutation and recombination. The coronavirus macrodomain, Mac1, is conserved amongst all known coronaviruses and is also conserved in the *Togaviridae* and *Hepeviridae* families. Mac1 is a key factor in viral replication and pathogenesis, but its role in the replication cycle remains unclear. A deeper investigation of Mac1 function will identify conserved antiviral mechanisms and aid in the development of Mac1 inhibitors that represent a novel strategy for antiviral therapeutics.

## INTRODUCTION

Coronaviruses (CoVs) are large, positive-sense ssRNA viruses with unusually large genomes ranging from 28-33 kb. Subdivided into four genera (alpha, beta, gamma, and delta), coronaviruses cause disease of clinical and veterinary significance in mammals and birds. Deadly infections in piglets with porcine endemic diarrhea virus illustrate the significant economic burden coronaviruses impose on the livestock industry (1); however, recent history demonstrates zoonotic transmission of CoVs to be an even greater economic and public health threat. Between 10-30% of circulating common cold viruses in humans are caused by CoVs like hCoV-229E and hCoV-OC43, which have a likely origin in zoonotic transmission from livestock(2, 3). The emergence of three highly pathogenic human CoVs within as many decades has placed the threat of novel zoonotic CoVs near the forefront of pandemic concern (4, 5). Efforts to determine the functions of highly conserved CoV proteins will help aid in developing novel therapies to help mitigate future outbreaks of divergent CoVs.

The CoV genome is organized into two major segments encoding the non-structural and the structural proteins. Structural proteins such as the spike and nucleocapsid protein are encoded in transcriptionally “nested” ORFs at the 3’ end of the genome. The non-structural proteins (nsps) are encoded at the 5’ end in two large co-terminal ORFs that comprise about two-thirds of the genome. Nsps are translated as large polyproteins from either ORF1a or from ORF1ab which are then proteolytically processed into each of the functional nsps. Translational regulation between these two ORFs is achieved through ribosomal frameshifting bypassing a conserved stop codon. It is thought this strategy is meant to maintain optimal stoichiometric ratios between proteins important for blocking immunity or establishing viral infection and the proteins that constitute the RNA replication machinery (6).

Nsp3 is the largest of the coronavirus nonstructural proteins with a mass of approximately 220 kDa. Nsp3 contains numerous domains with important characterized functions. These include polyprotein cleavage and deubiquitinase (DUB) activity of PL^pro^, ssRNA and nucleocapsid binding of Ubl1, ADP-ribosylhydrolase activity of the conserved macrodomain, and formation of replication organelles (ROs) (7–10). Most notably, nsp3-4 dimers form large hexamers that, on one side of the ER double membrane, complex with the replicase holoenzyme and, on the other side, comprise a pore through which newly synthesized viral RNA is shuttled into the cytoplasm(9, 11–13). These findings establish nsp3 as a platform for multiple key steps of the replication cycle, including the establishment of the RO, initial RNA replication, viral protein processing, and packaging. Interestingly, in SARS-CoV-2 infection, nsp3 has also been shown to be distributed at low levels throughout the cytoplasm, suggesting it may play additional roles in infection (14).

Within nsp3 reside up to three macrodomains termed Mac1, Mac2, and Mac3. Macrodomains are a class of conserved protein domains present in all forms of life. Their structure, characterized by a mixed α-β-α sandwich fold, is highly conserved, however their sequences – and therefore their functions – are diverse (15). Mac2 and Mac3, originally identified in SARS-CoV as part of the SARS-CoV Unique Domain (SUD), are present only in a subset of CoVs, most notably the *Sarbecoviruses*, and were demonstrated to bind nucleic acid (16). Mac1, on the other hand, is highly conserved across coronaviruses and has been shown to be a key virulence factor. Belonging to the MacroD-type class of macrodomains, the CoV Mac1 targets and hydrolyses ADP-ribose moieties, reversing a post-translational modification known as ADP-ribosylation (17–19).

ADP-ribosylation, the process by which ADP-ribose is covalently attached to target proteins, is facilitated by enzymes known as ADP-ribosyltransferases (ARTs), commonly referred to as PARPs. With 17 members in mammals, the PARP family regulates a myriad of cellular processes such as stress granule formation, cellular trafficking, DNA damage repair, gene expression, and cell cycle progression (20). Additionally, the importance of PARPs in the innate antiviral response is increasingly clear. Many of the mono-ADP-ribosylating (MARylating) PARPs are IFN-stimulated genes (ISGs) that modulate the inflammatory response and restrict viral replication (21–23). Importantly, ADP-ribose can be added to several different types of residues, including basic and acidic residues, as well as serine and cysteine (24, 25). Importantly, most macrodomains can only remove ADP-ribose from acidic residues but still bind to ADP-ribose attached at other residues. Thus, both ADP-ribose binding and hydrolysis can separately impact biological functions. In CoVs, Mac1 activity is required for IFN repression and efficient viral replication, though the exact targets remain unclear (26–31). Mac1 catalytic activity relies on a highly conserved asparagine (residue N1347 in mouse hepatitis virus (MHV) strain JHM (JHMV)) which coordinates critical hydrogen bonds with the distal ribose (32, 33). Previous characterizations of an alanine mutation at this site abolished the catalytic activity of Mac1 and sensitized the virus to type-I IFN (IFN-I) signaling but did not impact its ability to bind ADP-ribose (29, 34). In JHMV, a N1347A mutant virus replicated normally in IFN-deficient cells such as 17Cl-1 but was attenuated in IFN-competent bone-marrow derived macrophages (BMDMs). Importantly the replication defect was lost in IFNAR knockout cells, demonstrating that Mac1 catalytic activity counters IFN-mediated antiviral responses (34). Soon after, we also generated an alanine mutation at D1329 in JHMV, a residue predicted to coordinate stabilizing interactions with the adenine critical for substrate binding. This mutant demonstrated a large replication defect in both IFN-competent and IFN-deficient cells, suggesting differential roles for the binding and hydrolysis activity of Mac1 (29). Mutation in both residues resulted in a non-replicating virus that was unable to be launched *de novo.* Unsurprisingly, a full in-frame deletion of Mac1 from either JHMV or MERS-CoV resulted in unrecoverable viruses. Interestingly, however, SARS-CoV-2 tolerated the deletion, but was sensitive to IFN-γ pre-treatment, similar to the SARS-CoV-2 N1062A mutant virus (31). This result indicates that JHMV and MERS-CoV have a strict requirement for Mac1 ADP-ribose binding while SARS-CoV-2 may have evolved to encode for a protein with redundant function. We recently found that mutation of a conserved isoleucine in MERS-CoV and SARS-CoV-2 conferred enhanced Mac1 binding affinity to ADP-ribose with no impact on catalytic activity but resulted in an overall reduction in viral replication (35). This data together indicates that Mac1 is a critical factor in CoV replication and fitness during infection but relies on finely tuned biochemical properties to promote virus replication and pathogenesis.

In this study, we sought to define the stages of the viral replication cycle reliant on both known biochemical functions of Mac1, binding and catalysis. As predicted, we found that purified SARS-CoV-2 and MERS-CoV D-A mutant proteins demonstrated a stark decrease in substrate affinity which contrasted with the previously published N-A mutant protein. However, it retained substantial enzymatic activity as opposed to the N-A mutant. Thus, the D-A and N-A Mac1 mutants provide suitable models to better understand the role of ADP-ribose bindings versus hydrolysis in the viral lifecycle. In cell culture, the JHMV D1329A mutant virus showed a marked defect in RNA production, while the N1347A mutant virus resembled WT virus in RNA replication kinetics but was significantly reduced for protein accumulation in IFN-competent cells. These results provide a more refined understanding of the multiple mechanisms that Mac1 utilizes to promote CoV replication.

## RESULTS

### Mutation of conserved aspartic acid and asparagine residue to alanine dramatically reduces ADP-ribose binding but maintains ADP-ribosylhydrolase activity

Two highly conserved residues in JHMV Mac1, N1347 and D1329 **(Fig. 1A)** were previously demonstrated to impact viral fitness and virulence, though to different degrees and in a cell-type specific manner (29). To compare the biochemical properties of the D-A mutation with the previously characterized N-A mutation (29, 35), we performed several previously established ADP-ribose binding and hydrolysis assays using purified SARS-CoV-2 and MERS-CoV Mac1 WT and D-A proteins. We and others have been unable to produce MHV Mac1 protein, so SARS-CoV-2 and MERS-CoV proteins were used instead. ADP-ribose binding assays include a differential scanning fluorimetry (DSF) assay that measures thermal stability, a proximity-based AlphaScreen assay, and isothermal calorimetry (ITC) which measures the thermodynamic properties of binding. Thermal stability of the WT-Mac1 protein was demonstrated to be approximately 6°C higher than that of the D-A protein at the endpoint concentration of 1M ADP-ribose **(Fig. 1B)**. In the AlphaScreen assay, the D-A protein had a multi-log reduction in alpha counts compared to the WT protein **(Fig. 1C)**. Finally, isothermal titration calorimetry (ITC) data showed that the *K_D_* of Mac1 D-A to ADP-ribose was ∼16-fold higher than the WT-ADP-ribose interaction **(Fig. 1D)**. We also saw similar results for each of these assays using MERS-CoV Mac1 WT and D-A proteins **(Fig. S1)**. These data together demonstrate a marked defect in the Mac1 D-A protein’s ability to bind to ADP-ribose and distinguishes the D-A mutant from the N-A mutant, which was previously shown to bind ADP-ribose at near WT levels (35). In addition, we performed an *in vitro* ADP-ribosylhydrolase assay by co-incubating purified SARS-CoV-2 Mac1-WT and D-A proteins with mono-ADP-ribosylated (MARylated) PARP10 and measured the loss of MAR signal from PARP10 by Western blotting, as previously described (27). We found that Mac1-D-A retained substantial enzyme activity which contrasts with Mac1-N-A which we previously found to be largely devoid of catalytic activity **(Fig. 1E)** (35) These results are consistent with results from alphavirus macrodomain studies. These biochemical studies provide a stark delineation between the D-A and N-A mutant proteins as being deficient in either binding or hydrolysis, respectively.

**Fig. 1.**
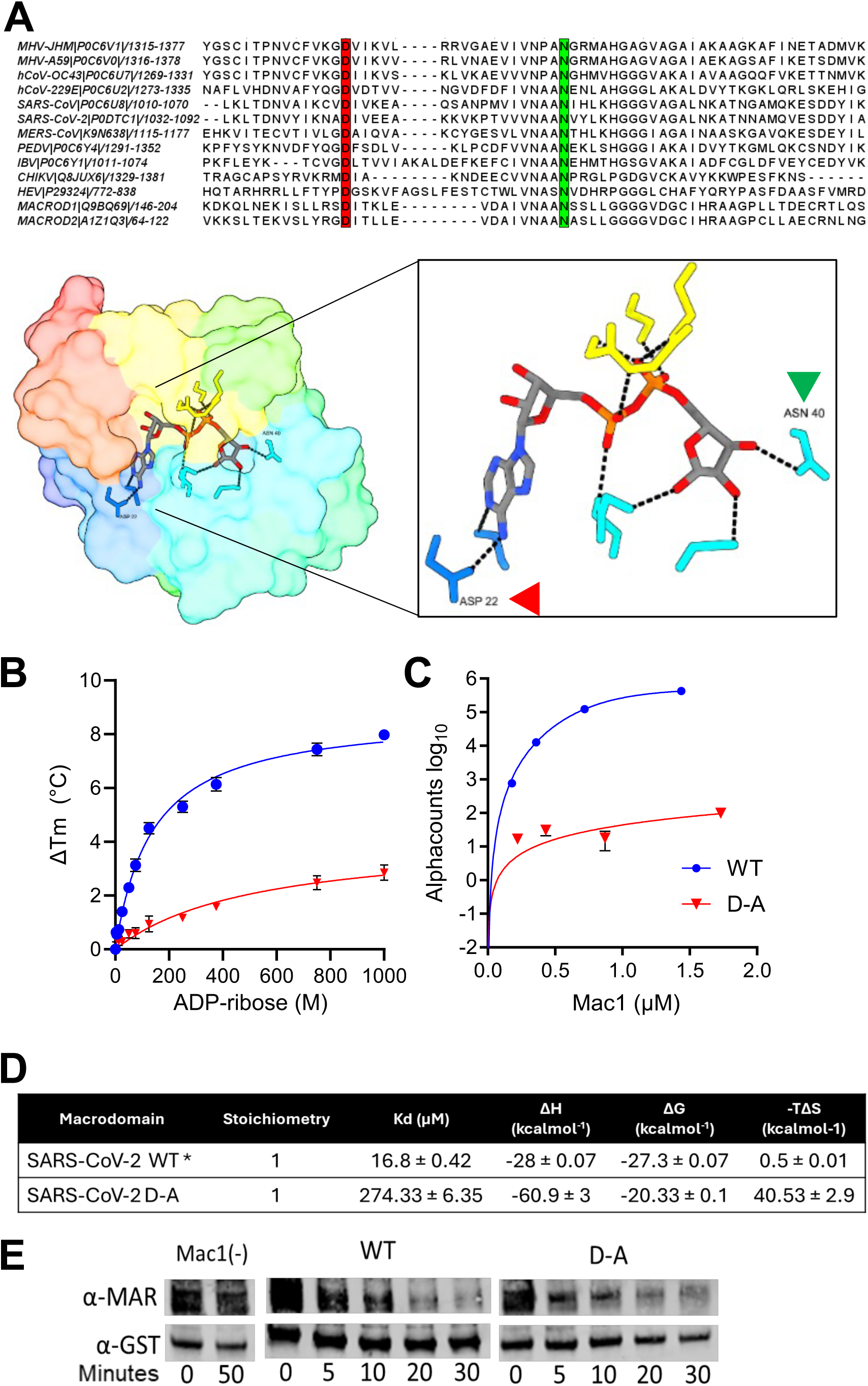
Mutation of a highly conserved aspartic acid in the ADP-ribose binding pocket of Mac1 dramatically reduces ADP-ribose binding but largely maintains its enzyme activity. **(A)** Highly conserved aspartic acid (red) and asparagine (green) residues highlighted in alignments of coronavirus Mac1 and other related macrodomain proteins. **(B)** SARS-CoV-2 Mac1-WT and D-A proteins were incubated with increasing quantities of ADP-ribose and then assayed for thermal stability by differential scanning fluorimetry (DSF). **(C)** SARS-CoV-2 Mac1 WT and D-A proteins were tested for their ability to interact with an ADP-ribosylated peptide using an AlphaScreen assay. The results in B-C are representative of 2 independent experiments. **(D)** Isothermal titration calorimetry (ITC) of purified SARS-CoV-2 WT and D-A proteins in the presence of increasing concentrations of ADP-ribose. *SARS-CoV-2 WT protein was previously published (27). The results are the combined averages of 4 independent experiments. **(E)** WT and D-A proteins were incubated with MARylated GST-PARP10 CD for the indicated time and MARylation levels were measured by Western blotting for mono-ADP-ribose. GST antibody was used as a loading control. Results are from one experiment representative of 2 independent experiments.

### MHV D1329A/D1330A virus is attenuated in cell culture and *in vivo*

Previous work with the JHMV single-nucleotide D1329A mutant virus revealed a high propensity for reversion *in vivo*, indicating a high fitness cost compared to other viable Mac1 mutants (29). To prevent this reversion, the JHMV-D1329A mutant virus was regenerated to contain a two-nucleotide substitution using a bacterial artificial chromosome (BAC) based reverse genetic system (36). We produced this virus in cell culture and confirmed that it had no reverting mutations. Furthermore, to confirm that the defect is not due to second-site mutations produced during cloning, we reintroduced the WT Mac1 sequence into the JHMV-D1329A BAC which was termed the “repaired” D1329A (*rep*D1329). In addition, we fully sequenced the MHV genome from terminal passages of D1329A and found no additional mutations in the viral genome. At low (0.1) and high (1.0) MOI, this new JHMV-D1329A mutant virus demonstrated the same multi-log replication defect in DBTs when compared to *rep*D1329 and N1347A viruses as previously described **(Fig. 2A-B)** (29). Importantly, intranasal infection of C57BL/6 with JHMV-D1329A resulted in no lethality or weight loss as compared to infection with *rep*D1329 which resulted in ∼25% mouse survival and an average of more than 10% weight loss. These results indicate that this two-base pair mutant D1329A virus, unlike the single-base pair mutant, was unable to revert to WT *in vivo* **(Fig. 2C-D)**. We also engineered the orthologous mutation into MHV-A59, D1330A. The A59-D1330A virus had a mild replication defect in cell culture **(Fig. 2E-G)** but replicated poorly *in vivo* compared to WT virus **(Fig. 2H**), indicating that Mac1 ADP-ribose binding is critical across multiple strains of MHV.

**Fig. 2.**
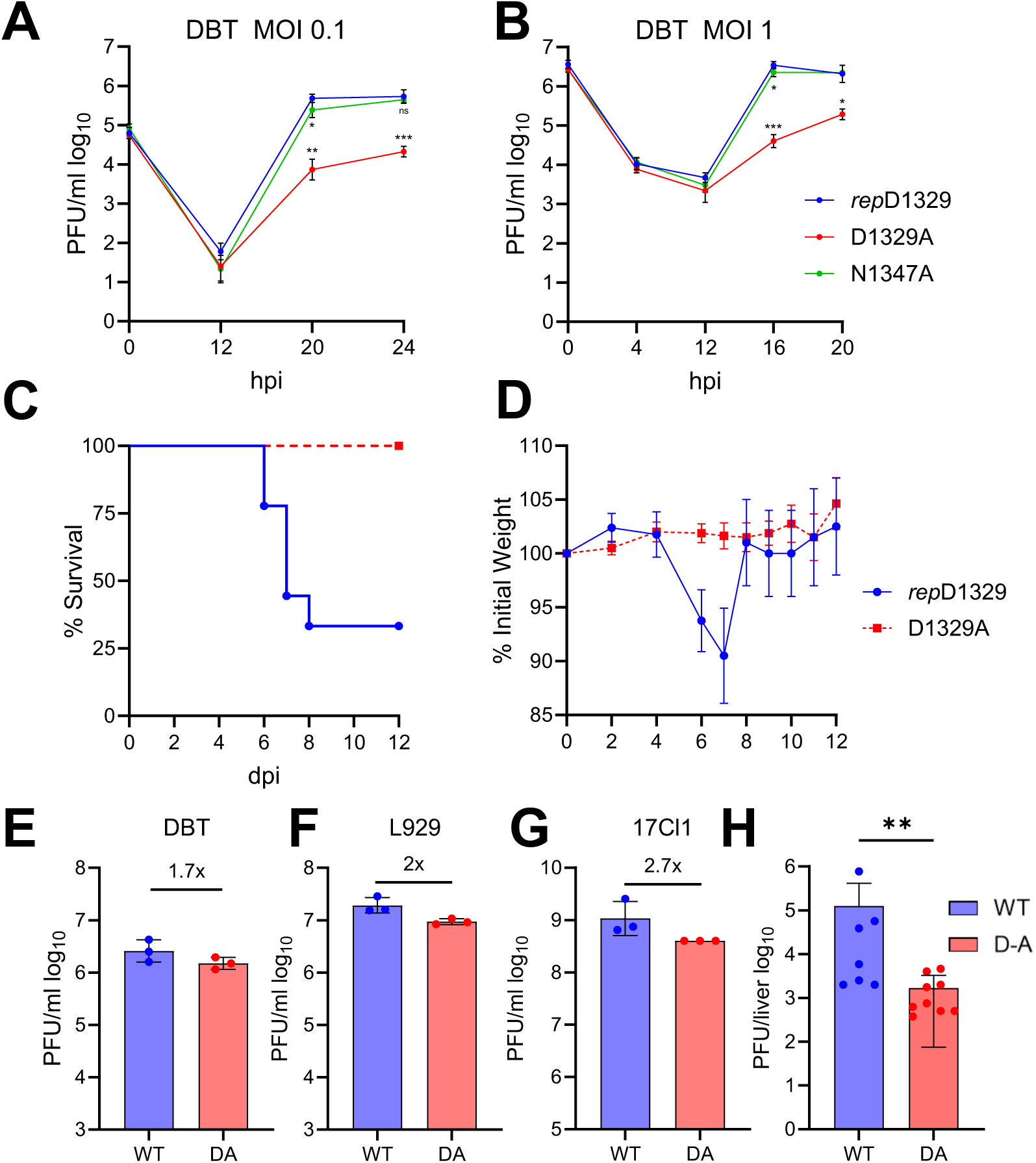
MHV-D1329A/D1330A virus is attenuated in cell culture and *in vivo.* **(A-B)** DBT cells were infected with JHMV-*rep*D1329, N1347A, and D1329A viruses at an MOI of 0.1 (A) and 1.0 (B). Progeny virus was collected at indicated times and was titered by plaque assay. The results are from a single experiment representative of 2 independent experiments. **(C-D)** C57BL/6 mice were infected with 10^4^ PFU JHMV-*rep*D1329 and D1329A virus and monitored for survival (C) and weight loss (D) over 12 days. n=9 mice, *rep*D1329; n=10 mice, D1329A. The results are the combined data from two independent experiments. **(E-G)** DBT (E), L929 (F), and 17Cl-1 (G) cells were infected with MHV-A59 WT and D1330A viruses and progeny virus was measured at 24 hpi. The results are from a single experiment representative of 2 independent experiments. **(H)** C57BL/6 mice were infected with 500 PFU A59-WT and D1330A viruses. Livers were collected at 3 dpi in PBS and viral loads were measured by plaque assay. Graphs represent the geometric mean with 95% confidence internal. n=7 mice, WT; n=9 mice, D1330A.

### JHMV-D1329A produces small plaques and demonstrates reduced infectivity in mouse cells compared to *rep*D1329 and N1347A viruses

The first notable hallmark of the JHMV-D1329A virus infection was a substantial reduction in overall plaque size in both DBT and L929 cells **(Fig. 3A-B)**. We quantitated the size of D1329A plaques as compared to *rep*D1329 and N1347A viruses and found that the D1329A plaques were on average about half the size of the plaques produced by *rep*D1329 and N1347A on both DBT and L929 cells **(Fig. 3C-D)**. JHMV plaques are formed by cell-cell syncytia mediated by the S protein (37). Thus, this phenotype is likely due to a severe reduction in the production of S protein or its ability to traffic to the cell surface. A severe lack of S protein production could be due to the inability to initiate infection in some cells. To determine if JHMV-D1329A has a defect in initiating a productive infection we performed a plaquing efficiency assay using DBT, L929, and HeLa cells expressing the MHV receptor (Hela-MVR). Equal plaque forming units (PFU) of *rep*D1329, D1329A, and N1347A were plated on each cell type and the number of plaques were determined. D1329A produced ∼5-10-fold fewer plaques on both DBT and L929 cells as compared to *rep*D1329 and N1347A, indicating that D1329A has a substantial defect in the ability to initiate a productive infection. Interestingly, this defect may be species specific, as no defect was observed in Hela-MVR cells which are of human origin but express the MHV receptor **(Fig. 3E)**.

**Fig. 3.**
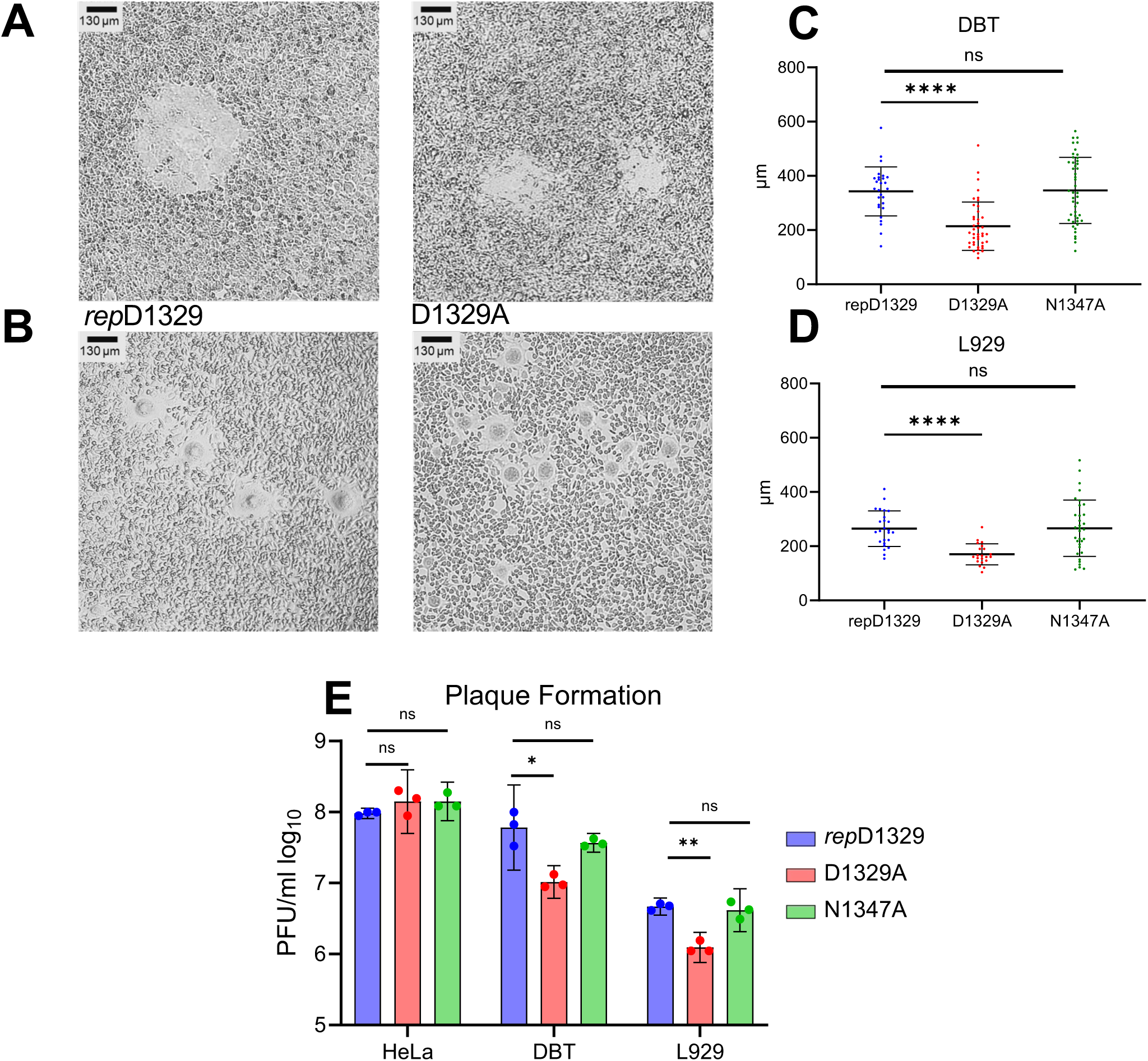
JHMV-D1329A virus produces small plaques and demonstrates reduced infectivity in mouse cells compared to *rep*D1329 and N1347A viruses. **(A-B)** DBT (A) and L929 (B) cells were infected with JHMV-*rep*D1329 and D1329A at an MOI of 0.1 and were examined at 18 (DBT) or 20 (L929) hpi to determine relative plaque sizes between mutants. **(C)** Plaque sizes were measured using CaptaVision software calibrated with a stage micrometer n=27, *rep*D1329; n=41, D1329A; n=45, N1347A **(D)** Plaque sizes were measured using CaptaVision software calibrated with a stage micrometer n=25, *rep*D1329; n=21, D1329A; n=33, N1347A The results represent the combined results of 2 independent experiments. **(E)** Hela-MVR, DBT, and L929 cells were infected with equal PFUs of each virus and the number of plaques were counted 20 hpi. The results are from a single experiment representative of 3 independent experiments.

### JHMV-D1329A has no defect in genomic RNA delivery into cells

To determine if the inability of D1329A virus to initiate infection stems from a reduction in the entry of viral RNA into the cytoplasm of infected cells, we measured intracellular genomic RNA levels at 2 hpi via RT-qPCR. Each of the three indicated cell lines were infected with JHMV-*rep*D1329 and D1329A for two hours; after which, the cells were trypsinized and washed three times with PBS to remove non-infective virions then resuspended in TRIzol™ Reagent (Invitrogen). RT-qPCR revealed no significant reduction in D1329A gRNA levels when compared with *rep*D1329, suggesting that the plaquing defect is not due to a defect in viral entry **(Fig. 4)**.

**Fig. 4.**
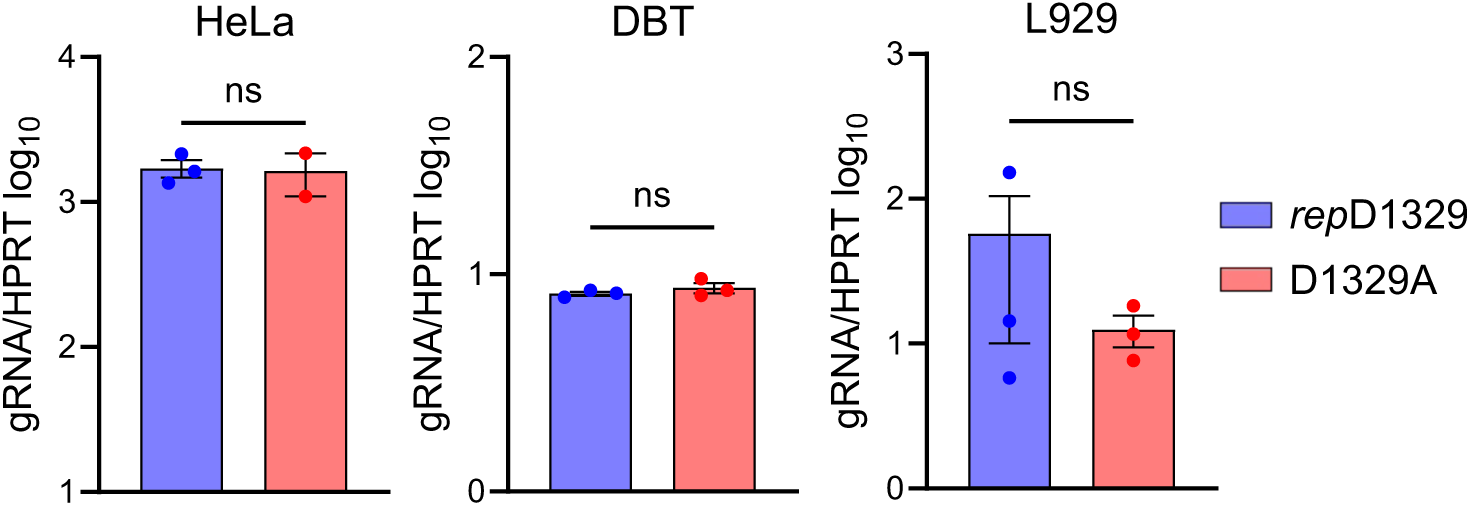
JHMV-D1329A virus has no defect in genomic RNA delivery into cells. Indicated cell lines were infected with JHMV *rep*D1329 and D1329A viruses for 1 hr, then trypsinized for 5 minutes and rinsed 3 times with PBS. The cells were then collected, RNA was harvested, and genomic RNA levels were determined by qPCR using the *Δ*Ct method and normalized to HPRT mRNA. These data are from a single experiment representative of 2 independent experiments.

### JHMV-D1329A virus is defective in genomic and sub-genomic RNA accumulation compared to *rep*D1329 and N1347A viruses

Next, we assayed the accumulation of viral genomic and sub-genomic RNAs in both high and low MOI infections of DBT and L929 cells. Using RT-qPCR, viral genomic-RNA (gRNA) and sub-genomic RNA6 (sgRNA6) levels were quantified over a 20-hour period **(Fig. 5A-B)**. We used sgRNA6 as a marker for sub-genomic RNAs as the primers for sgRNA6 gave the most robust results in our qPCR assay. At high MOI, the N1347A virus closely resembled the *rep*D1329 virus, as viral sgRNA6 and gRNA levels increased substantially between 4 and 12 hpi in DBTs, while D1329A infected cells failed to produce any new viral transcripts before 12hpi. This difference was most notable in sgRNA6, as *rep*D1329 and N1347A transcripts were increased by nearly 3-logs over the D1329A virus. The D1329A infected cells did show increasing RNA production after 12hpi, indicating a delay in transcription initiation, though the levels never reached that of *rep*D1329 or N1347A infected cells **(Fig. 5A)**. Similarly, infection of L929s showed approximately 10 to 100-fold increases in gRNA and sgRNA6 levels in the *rep*D1329 and N1347A infected cells compared to D1329A throughout the course of infection **(Fig. 5B)**. Similar results were seen at low MOI **(Fig. S2)**. To determine whether the defect in D1329A RNA accumulation was due to targeted degradation of viral transcripts, infected L929 cells were treated with the Remdesivir metabolite GS-441524 at 10 hpi to arrest RdRp-mediated transcription (38), after which viral transcripts were quantified at four-hour intervals until 22hpi **(Fig. 5C)**. GS-441524 was not added until 10 hpi as earlier addition may have limited the amount of RNA produced during D1329A infection making it unfeasible to detect RNA degradation. Viral RNA synthesis increased in GS-441524 treated cells for a few hours, after which they stabilized, as compared to RNA levels in untreated cells, which continued to increase. In both the *rep*D1329 and D1329A infections treated with GS-441524, there was no evidence of RNA degradation as RNA levels did not significantly decrease through the course of the infection, indicating that there is no increased degradation of the D1329A transcripts compared to *rep*D1329.

**Fig. 5.**
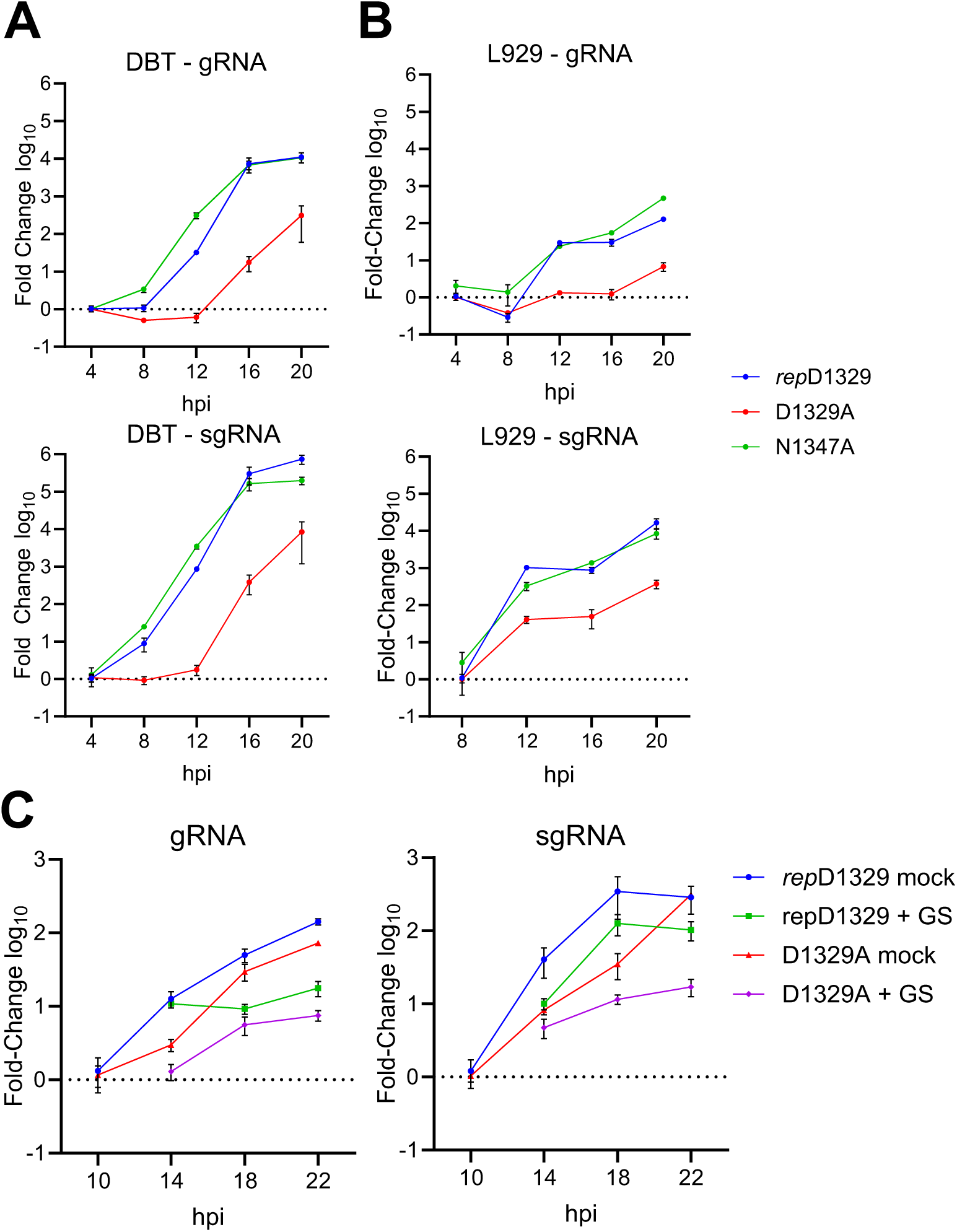
JHMV-D1329A virus is defective in the production of viral genomic RNA (gRNA) and sub-genomic RNA6 (sgRNA6) compared to *rep*D1329 and N1347A viruses. **(A-B)** DBT (A) and L929 (B) cells were infected with indicated viruses at an MOI of 3.0. Cells were collected at indicated times and gRNA (top) and sgRNA6 (bottom) were measured by qPCR using the *ΔΔ*CT method and normalized to HPRT. The data in A-B is from a single experiment representative of 2 independent experiments. **(C)** L929 cells were infected with either *rep*D1329 or D1329A viruses at an MOI of 0.1. At 10 hpi cells were treated with Remdesivir metabolite GS-441524 or DMSO control. Cells were collected at indicated time points and viral gRNA (left) or sgRNA6 (right) levels were determined by qPCR using the *ΔΔ*Ct method and normalized to HPRT mRNA. The results are from a single experiment representative of 2 independent experiments.

As a separate method to analyze viral RNA production, we performed confocal immuno-fluorescence microscopy with probes for dsRNA, the hallmark RNA replication intermediate **(Fig. 6A)**. DBT cells were infected with *rep*D1329, D1329A, and N1347A at an MOI of 0.5 and cells were fixed and stained for nsp3 and dsRNA at 8 hpi. While mock infected cells were devoid dsRNA signal, *rep*D1329 and N1347A infected cells demonstrated robust dsRNA staining that co-localized with nsp3, indicating the dsRNA represented viral RNAs. In contrast, D1329A infected cells had minimal dsRNA despite having significant nsp3 staining. Quantitation of these images confirmed that the D1329A virus produced significantly reduced levels of dsRNA but only mildly reduced levels of nsp3 **(Fig. 6B)**. In addition, we measured the proportion of dsRNA that overlapped with nsp3 (labeld as M1) and conversely the proportion of nsp3 that overlapped with dsRNA (labeled as M2) in each image using a Mander’s coefficient. We found that wherever dsRNA staining was present, it almost always overlapped with nsp3 staining from each virus **(Fig. 6C, M1-green)**. However, there were a couple of cases in D1329A infected cells where nsp3 staining did not contain any overlapping dsRNA staining, unlike *repD*1329 or N1347A **(Fig. 6C, M2-red)**. This result indicates that D1329A infected cells produced nsp3 but were unable to efficiently produce RNA. N1347A, conversely, appeared to produce similar levels of nsp3 and dsRNA staining compared to *rep*D1329. To further determine if D1329A can efficiently translate non-structural proteins, we stained for nsp3 at 6 hpi, the earliest time point that we could detect it. Imaging demonstrated that D1329A produced nsp3 at these early time points at levels that were reduced but not significantly different from *rep*D1329 and N1347A **(Fig. S3A-B)**. The reduction in nsp3 is likely due to already reduced RNA production in these cells, as more genomic RNA would lead to increased production of nsps for the *rep*D1329 or N1347A viruses. In combination, these results suggest that Mac1 ADP-ribose binding is required for efficient viral transcription initiation in murine cells, whereas hydrolysis activity is likely dispensable in the earliest stages of replication. **N1347A infected primary bone-marrow derived macrophages (BMDMs) produce WT levels of viral RNA but have reduced N protein accumulation compared to *rep*D1329 virus.** We previously demonstrated that both N1347A and D1329A mutants have similar replication defects in primary macrophages when infected at low MOI, which could be partially rescued by treatment with PARP inhibitors (29). However, N1347A replication could also be rescued in IFNAR and PARP12 knockout cells, while D1329A replication was not, indicating these viruses may impact different parts of the viral lifecycle in these cells (31, 34). First, we tested whether these viruses had replication defects at high MOI so we could evaluate the viral lifecycle within a single cycle of replication. As expected, both viruses had replication defects in BMDMs when infected at an MOI of 1.5 **(Fig.7A)**. Specifically, D1329A virus titers were reduced almost 14-fold from *rep*D1329, while N1347A titers were reduced 5-fold. Next, we evaluated the accumulation of viral gRNA and sgRNA6. D1329A infection resulted in a significant reduction in both gRNA and sgRNA6 levels at 9 hpi, while gRNA and sgRNA6 levels following N1374A infection were similar to those of *rep*D1329, as seen in both DBT and L929 cells **(Fig. 7B)**. Specifically, sgRNA and gRNA levels were mildly reduced at 9 hpi but increased at 12 hpi following N1347A infection, though these differences were not significant. However, when we evaluated the production of viral structural proteins, N1347A infected cells produced reduced levels of nucleocapsid (N) and spike (S) proteins at 9 and 12 hpi **(Fig. 7C)**. When quantitating viral protein levels from multiple experiments, the amount of N protein was significantly reduced almost 4-fold at 9 hpi compared to *rep*D1329 infection, while at 12 hpi N protein was reduced 1.5-fold **(Fig. 7D)**.

**Fig. 6.**
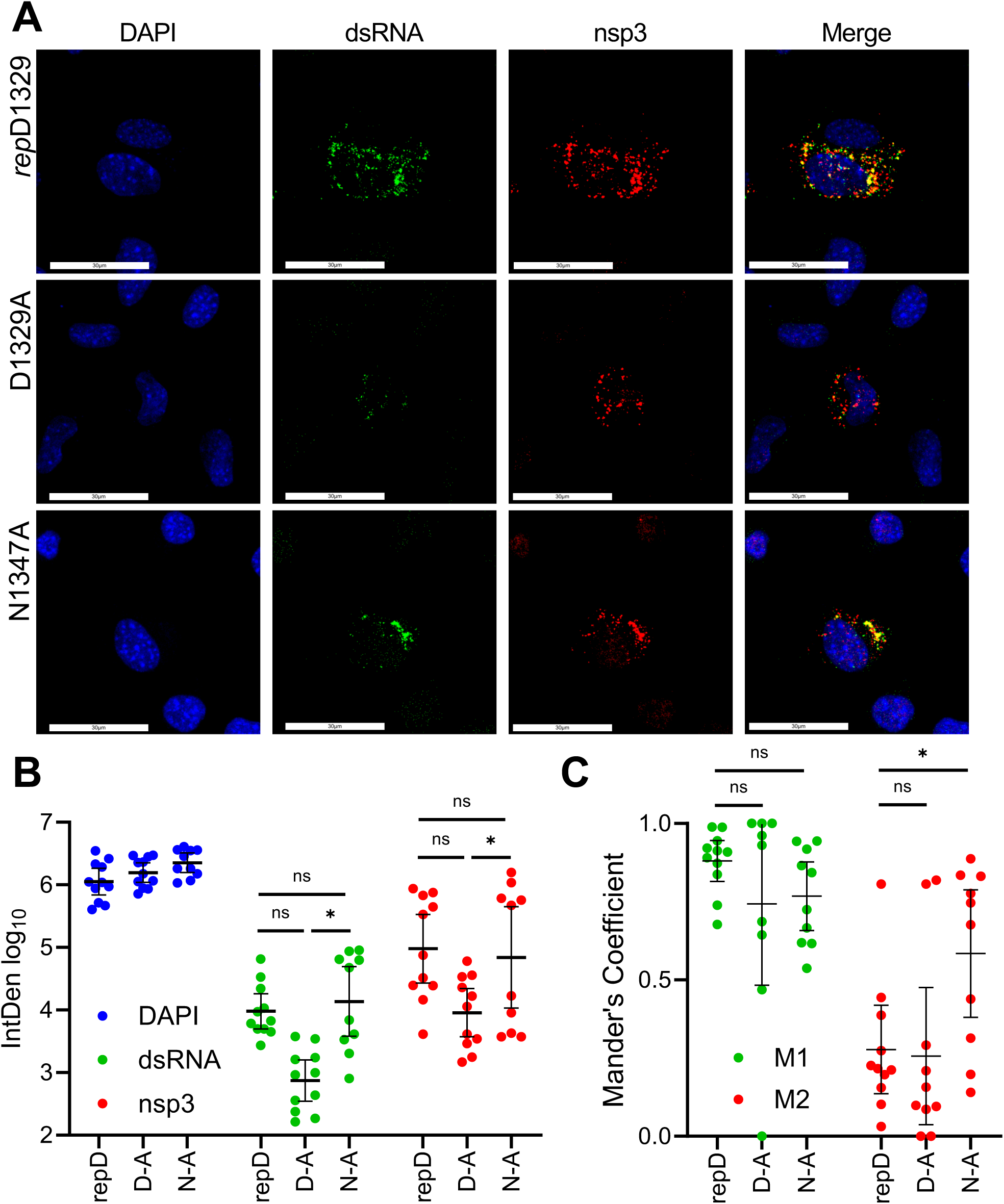
JHMV-D1329A virus has reduced dsRNA levels early in infection. **(A)** DBTs were infected with indicated virus at an MOI of 0.5, fixed at 8 hpi, stained for dsRNA and nsp3, and analyzed by confocal microscopy. **(B)** Quantification of fluorescence signal displayed as a product of mean pixel intensity and thresholded area (IntDen). Error bars indicate geometric mean with 95% CI, combined data from two separate experiments. **(C)** Mander’s coefficients displaying the proportional overlap between regions where dsRNA overlaps nsp3 staining (M1 – green), and where nsp3 staining overlaps dsRNA (M2 – red). Error bars indicate mean with 95% CI, combined data from two separate experiments.

**Fig. 7.**
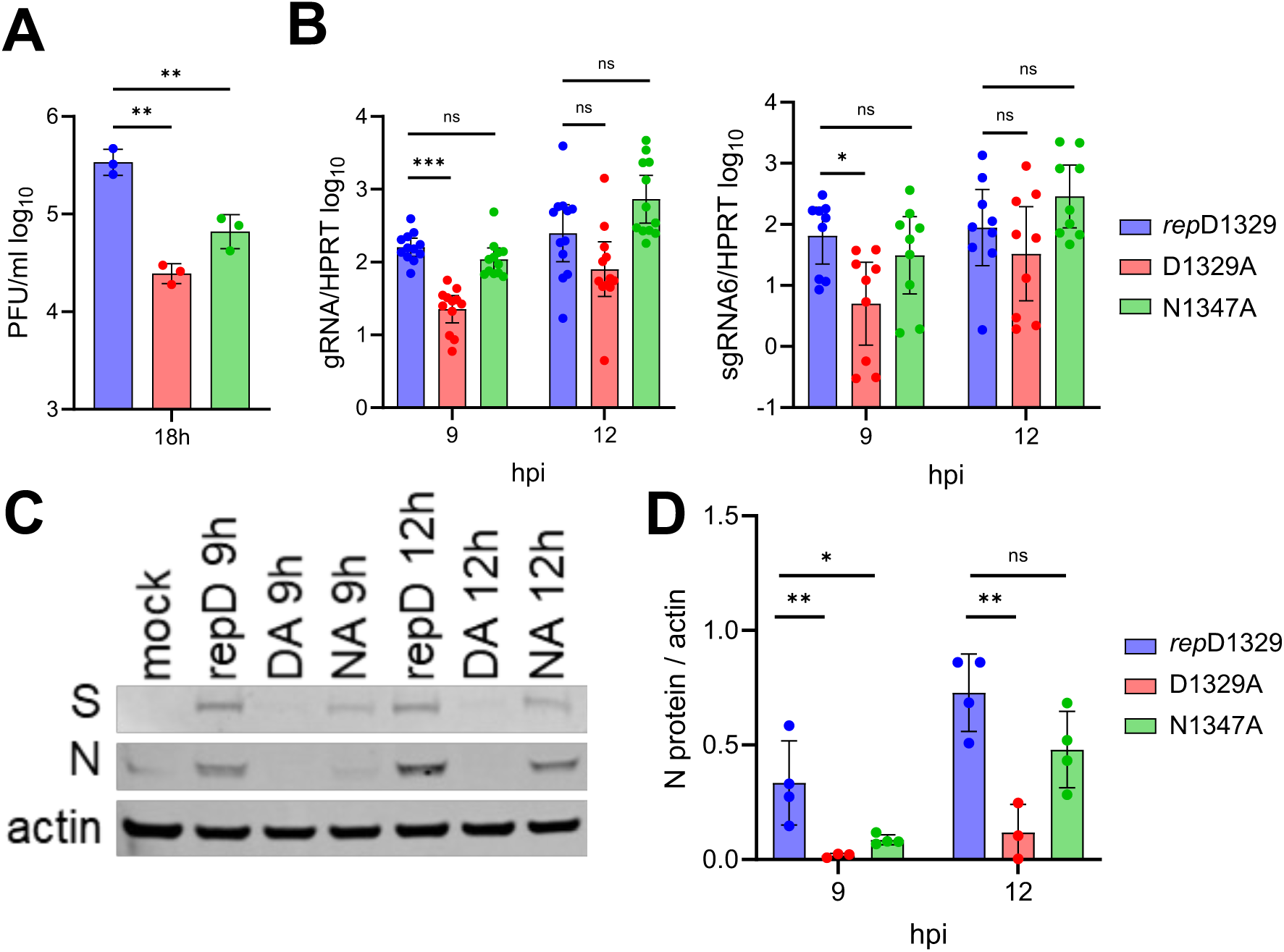
N1347A infected BMDMs produce WT levels of viral RNA but have reduced N protein accumulation compared to *rep*D1329 virus. **(A)** M2 differentiated BMDMs were infected with indicated viruses at an MOI of 1.5. Cells and supernatants were collected at 18 hpi and progeny virus was measured by plaque assay. The results are from a single experiment representative of 2 independent experiments. **(B)** BMDMs were infected as in (A) and cells were collected at indicated times points and gRNA (left) and sgRNA6 (right) were measured by qPCR using the *Δ*Ct method and normalized to HPRT mRNA. The data in B represent the combined results of 3 (sgRNA6) or 4 (gRNA) independent experiments. **(C)** BMDMs were infected as in (A) and cells were collected in sample buffer at indicated times points and viral spike (S) and nucleocapsid (N) proteins were analyzed by Western blotting. Actin was used as a loading control. The results are from a single experiment representative of 2-4 independent experiments. **(D)** Quantitation of the western blot signal in C normalized to actin using Image J. The data in (D) represent the combined results from 4 (*rep*D1239 & N1347A) or 3 (D1329A) independent experiments.

Additionally, we evaluated nsp3 and N protein abundance by confocal immunofluorescence in BMDMs. BMDMs were infected at an MOI of 0.5 before fixation and staining for nsp3 and N protein at 9 hpi **(Fig. 8A)**. Nsp3 puncta were again observed across all three viruses. Quantification showed the D1329A infected cells had 7.5-fold less nsp3 staining compared to *rep*D1329, while nsp3 levels were only reduced 1.6-fold in N1347A infected cells, neither of which were significant **(Fig. 8B, middle)**. In fact, in several cells infected with D1329A, the level of nsp3 was at or just below the mean of the *rep*D1329 infected cells, again demonstrating that D1329A can produce sufficient levels of nsp3 early in infection. This is in contrast to N protein staining, which was reduced nearly 100-fold in D1329A infected cells and almost 10-fold in N1347A infected cells **(Fig. 8B, right)**. However, the level of N protein staining in N1347A infected cells was highly variable within individual cells, with some cells producing normal levels of N protein, while others were dramatically reduced. As N1347A is restricted by IFN-induced PARPs (34), these results suggest that the levels of PARP induction between individual cells may fluctuate, resulting in inconsistent reductions in N protein production compared to *rep*D1329. But in general, N1347A infected cells produced less N protein, but not nsp3, than the *rep*D1329 infected cells. These observations suggest that initial nsp3 synthesis is not substantially impacted by Mac1, but that subsequent viral RNA or late protein production relies on the binding and hydrolysis activities, respectively, of Mac1 to proceed efficiently.

**Fig. 8.**
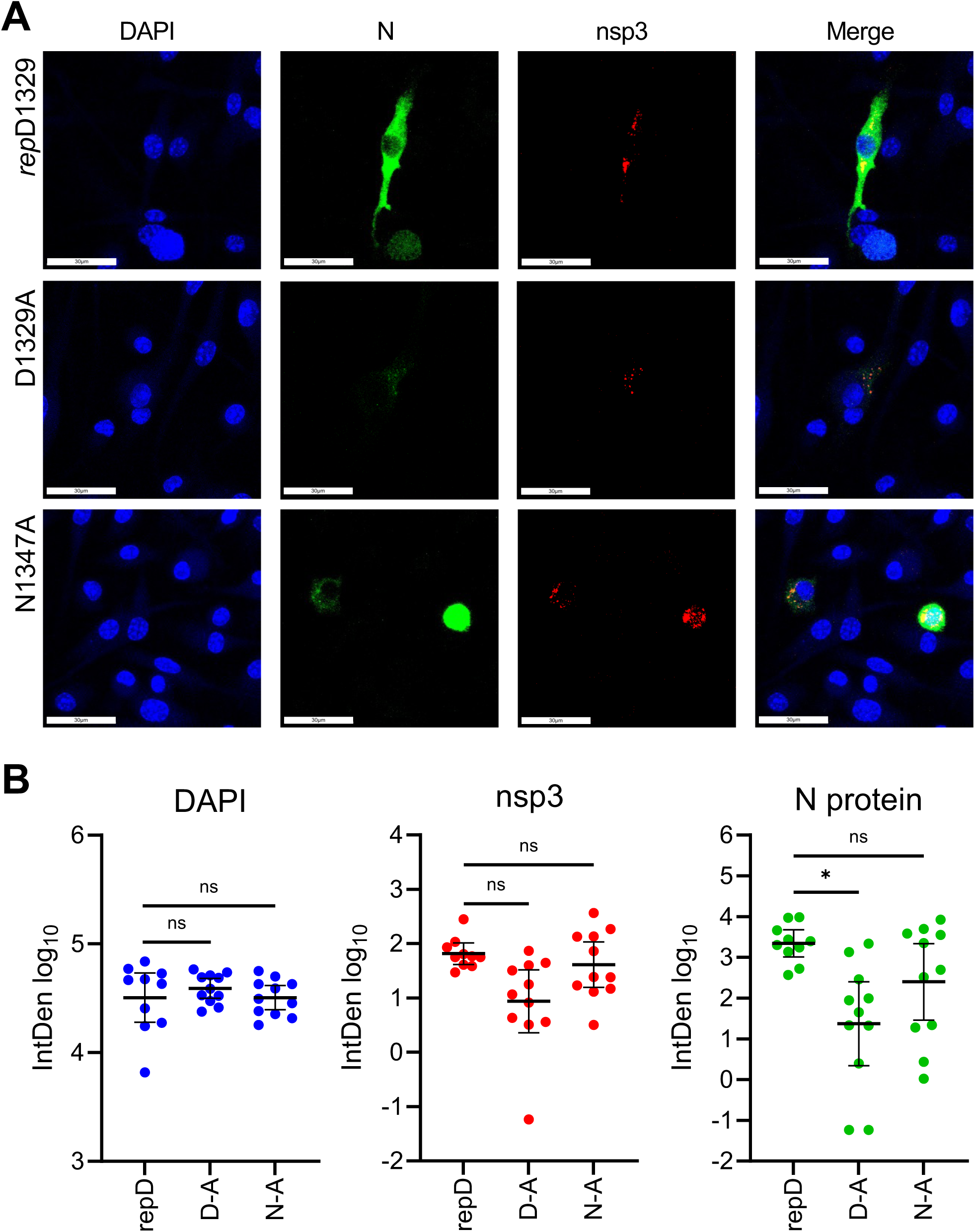
D1329A and N1347A infected BMDMs have near *rep*D1329 levels of nsp3 but have reduced N protein abundance. **(A)** BMDMs were infected with indicated virus at an MOI of 0.5 and fixed at 9 hpi, stained for nsp3 and N protein, and analyzed by confocal microscopy. **(B)** Quantification of fluorescence signal displayed as a product of mean pixel intensity and thresholded area (IntDen) for all three fluorescent channels. Combined data from two independent experiments presented as geometric mean with 95% confidence intervals.

## DISCUSSION

The current understanding of the conserved CoV macrodomain relies largely on its characterization as an ADP-ribosylhydrolase. Previous work has highlighted the importance of PARP activity and Mac1’s de-ADP-ribosylating activity on viral fitness and pathogenesis in both coronaviruses and alphaviruses (39). We have previously generated alanine mutations in several conserved residues of the Mac1 binding pocket. We found that mutation in a conserved asparagine residue important for hydrolysis activity, N1347A in JHMV, attenuated the virus in primary macrophages and sensitized it to PARP activity. We also found that mutating another conserved residue predicted to coordinate binding affinity, D1329, conferred a greater replication defect for this virus. Finally, we have been unable to recover JHMV that incorporates both mutations or a full Mac1 deletion. These findings led us to hypothesize that Mac1 is critical for JHMV infection and serves multiple roles in the viral replication cycle.

In this study, we sought to determine the stages of the viral life cycle most reliant on the binding and hydrolysis activity of Mac1 in an effort to better understand the roles Mac1 plays in infection. To characterize the biochemical impacts of the conserved aspartic acid and asparagine residues in the ADP-ribose binding pocket, we utilized purified SARS-CoV-2 and MERS-CoV Mac1 proteins with each mutated to alanine and then performed *in vitro* ADP-ribose binding and hydrolysis assays. These data demonstrated a stark delineation between these mutant Mac1 proteins and their ADP-ribose binding and hydrolysis activities. Previous work from our group established the N-A mutant Mac1 suffered little to no loss in substrate binding affinity but had greatly reduced hydrolysis (35) while here we found that the D-A mutant Mac1 had a multi-log reduction in binding affinity, but only a slight reduction in hydrolysis. Importantly, similar results were demonstrated by other groups with SARS-CoV-2 mutant proteins and with recombinant alphavirus macrodomain proteins as well (33, 40). Next, we sought to compare the impact of both mutations on viral replication in the early viral life cycle. To achieve this, we utilized the previously generated JHMV-N1347A virus and re-generated a new clone of the previously studied D1329A virus. This new clone was generated via a two base-pair substitution in the D1329 codon rather than one to reduce the likelihood of reversion. As expected, this virus demonstrated severe attenuation in cell culture and *in vivo*. Through the detection of dsRNA by confocal microscopy and specific determination of viral genomic and sub-genomic RNA levels by qPCR, we found that the D1329A virus does not efficiently initiate viral RNA production following infection of any murine cell line that is susceptible to JHMV **(Figs. 5-6)**. Furthermore, the D1329A virus produced similar amounts of nsp3 in BMDMs as *rep*D1329 virus, and puncta of nsp3 staining in D1329A infected cells often lacked any notable dsRNA staining. In contrast, the N1347A virus replicated viral RNA at levels equal to that of WT virus in all cell lines tested but had a notable defect in viral protein production in primary BMDMs **(Fig. 7)**. These data indicate that the binding activity of Mac1 is key for early RNA production, while the hydrolysis activity is important for accumulation of viral proteins later on in the replication cycle. However, we cannot rule out the possibility that other defects, such as in polyprotein processing or assembly, may also be present. In fact, considering the level of N protein in N1347A infected BMDMs was almost back to WT levels at 12 hpi, it seems likely that there may be further defects in the assembly process for this virus. Below we present several hypotheses for how each activity may promote replication.

### ADP-ribosylhydrolase activity

We have demonstrated that PARP12 and PARP14 are both required to restrict the replication of CoVs with catalytically inactive Mac1 (21, 31, 41). Others have found that PARP14 is required for the production of ADP-ribose puncta following IFN*γ* or poly(I:C) treatment (34). Within these puncta, both PARP14 and DTX3L, an E3 ubiquitin ligase, are likely ADP-ribosylated, with DTX3L/PARP9 regulating the activity of PARP14 (42, 43). As an E3 ubiquitin ligase, it’s possible that DTX3L ADP-ribosylation enhances its ability to target viral proteins for degradation. However, there are many other potential targets of PARP14, such as p62, an autophagy regulator, and RACK1, a ribosome binding protein, that could impact viral protein production through autophagic degradation or repression of translation, respectively (44, 45). Furthermore, PARP12 is localized to the Golgi and cell stress can trigger PARP12 movement from the Golgi and a block in anterograde-membrane trafficking, which could impact viral assembly (46). Further work is required to decipher which PARP12/14 target(s) restrict viral protein production or assembly following infection with Mac1 catalytic-mutant CoVs.

### ADP-ribose binding activity

We envision two separate but not mutually exclusive hypotheses for how this Mac1 function promotes viral RNA production. Recent work has demonstrated that nsp3 plays a major role in forming a hexameric complex that constitutes a pore in the replication organelle, through which viral RNA is thought to pass (9, 11). As part of the larger pore structure, the N-terminal domains of nsp3 extend into the cytoplasm, where one or more subunits may interact with a number of host and viral factors. Previous work has established the association of nsp3 with the nucleocapsid protein as a critical factor in establishing CoV infection (8, 47, 48). Current models postulate that, since CoV gRNA serves as both the initial mRNA for nsp translation and the template for RNA replication, N protein may shuttle gRNA into the replication organelle via the nsp3 pore (8). Mutational analysis from several groups have demonstrated this relies largely on the interaction of the N-terminal Ubl1 of nsp3 with multiple phosphorylated residues in the serine and arginine-rich regions N1b and N2a of N protein (47–49). Other characterizations of the CoV nucleocapsid have revealed that it is also ADP-ribosylated (50). From the current data, it appears that the N protein is ADP-ribosylated only during infection, and the modification is retained in the virion. Moreover, the modification was shown to be unaffected by catalytically active Mac1. Thus, in our first hypothetical model, Mac1 would interact with an ADP-ribosylated N protein, to promote the translocation of either the viral genome or N protein through the pore **(Fig. 9A)**. Since N protein phosphorylation is reliant on host kinases (51) and ADP-ribosylation may compete with phosphorylation on some residues, perhaps Mac1 binding to ADP-ribosylated N evolved as an alternative measure to maintain the interaction between N protein and nsp3. Identification of the sites of N protein ADP-ribosylation and mutation analysis would help clarify how ADP-ribosylation affects N protein’s ability to interact with nsp3.

**Fig. 9.**
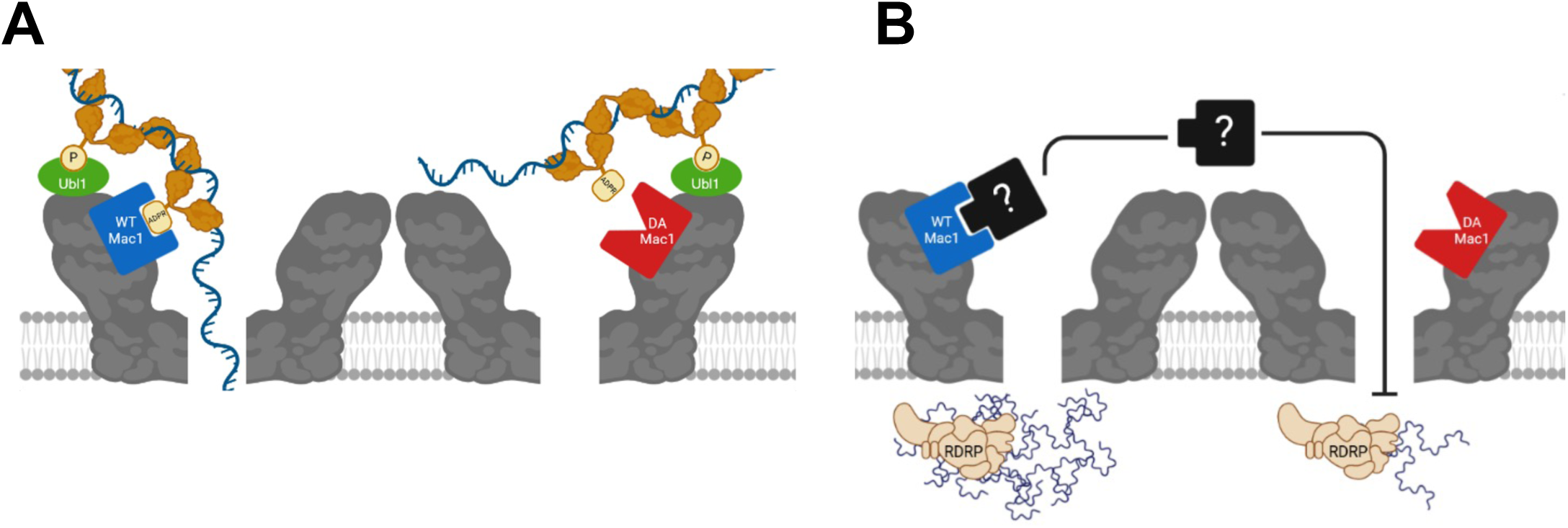
Potential roles for Mac1 in regulating early RNA transcription. **(A)** The Ubl1 domain of nsp3 interacts with phosphorylated residues on the RNA-bound nucleocapsid protein which is a key factor in shuttling viral RNA through the nsp3 pore to the site of transcription. Mac1 binding affinity may enhance this interaction between nsp3 and N protein, without which there is a delay in transcription initiation. **(B)** Mac1 binding affinity may sequester or block ADP-ribosylated host factors that target and inhibit viral transcription early in the replication cycle.

However, the hypothesis best supported by the current data is that the ADP-ribose binding affinity of Mac1 serves to prevent a key antiviral host factor from disrupting the initiation of viral transcription. Plaquing efficiency assays from this study showed that the D1329A mutant virus more readily establishes productive infections in HeLa-MVR cells than in murine DBT or L929 cells, suggesting a host-dependent anti-viral response. Furthermore, treatment with PARP inhibitors partially rescues D1329A virus replication, and treatment with the PARP substrate NAD^+^further reduces the replication of the D1329A virus (29). These results in combination suggests there is a murine-specific host factor that restricts virus replication in an ADP-ribosylation dependent manner. Binding of Mac1 to an unknown ADP-ribosylated host factor or factors may prevent it from traversing the nsp3 pore to access the replication transcription complex (RTC) and repress viral RNA production **(Fig. 9B)**. Structural analysis of the pore complex predicts a central channel of around 17 Å (52) that may be large enough for small proteins to access the RTC. Previous studies of related alphavirus macrodomain mutant viruses found that macrodomain/ADP-ribose binding is also required for initiating efficient viral RNA synthesis, indicating that there may be a conserved host mechanism that represses the initial transcription of multiple viruses (40, 53). Thus, future research into these mechanisms could identify novel methods for developing anti-viral therapeutics targeting multiple classes of RNA viruses.

## METHODS

### Cell Culture

Delayed Brain Tumor (DBT), L929, 17CL-1, and HeLa cells expressing the MHV receptor CEACAM1 (HeLa-MVR) were cultured in Dulbecco’s modified Eagle medium (Corning) supplemented with 10% fetal bovine serum, 100 U/ml penicillin, 100 mg/ml streptomycin, 10mM HEPES, 1mM sodium pyruvate, 0.1mM nonessential amino acids, and 2mM L-glutamine. Bone marrow-derived macrophages (BMDMs) were sourced from WT C57BL/6 mice and differentiated to M0 via culturing for 7 days in Roswell Park Memorial Institute medium (RPMI) supplemented with 10μg/ml Macrophage Colony Stimulating Factor (MCSF) (Genscript) in addition to 10% fetal bovine serum 100 U/ml penicillin, 100 mg/ml streptomycin, 10mM HEPES, 1mM sodium pyruvate, 0.1mM nonessential amino acids, and 2mM L-glutamine. Differentiation into M2 macrophages was conducted was achieved by the further addition of 10μg/ml IL-4 (Peprotech Inc.) preceding infection by 24 hours.

### Mice

Pathogen-free C57BL/6 mice originally sourced from Jackson Laboratories were bred and maintained in the animal care facility at the University of Kansas as approved by the University of Kansas Institutional Animal Care and Use Committee (IACUC) following guidelines set forth in the *Guidelines for the Care and Use of Laboratory Animals*.

### Generation of Recombinant JHMV and A59 Constructs

Recombinant pBAC-JHMV D1329A constructs were generated using Red recombination. Double-base pair mutations in the nsp3 Mac1 were engineered using the Kan^r^-I-SceI marker cassette as previously described, using primers listed in **Table 1**. Purified BAC DNA from Cml^r^ Kan^s^ colonies were analyzed by restriction enzyme digest, PCR and full plasmid sequencing for validation of correct clones. Primers used for validation of correct clones are also listed in **Table 1**. The recombinant MHV-A59 D1330A mutant was generated as described previously (54, 55). The D1330A mutation was introduced using site-direct mutagenesis (Q5 Site-Directed Mutagenesis Kit, E0554S, NEB England Biolabs) and sequencing-validated. The full-length viral cDNA was assembled using T4 DNA ligase (M0202L, NEB England Biolabs). The full-length viral RNA was synthesized using an in vitro transcription kit (mMESSAGE mMACHINE T7 Transcription Kit, AM1344, Thermo Fisher Scientific)

**TABLE 1.**
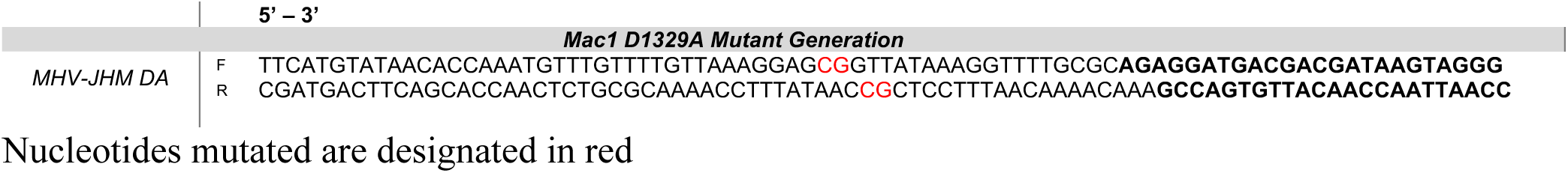
Linear recombination primers for engineering JHMV-D1329A.

### Reconstitution of recombinant pBAC-JHMV-derived virus

Approximately 1×10^6^ BHK-MVR cells were transfected with approximately 0.5-1 μg of pBAC-JHMV DNA and 1 μg of pcDNA-MHV-N plasmid or 1 μg of full-length A549 RNA using Polyjet (SignaGen) as a transfection reagent. Virus stocks were created by infecting ∼1.5×10^7^ 17Cl-1 cells at an MOI of 0.1 plaque-forming units (PFU)/cell and collecting both the cells and supernatant at 16-20 hpi. The cells were freeze-thawed, and debris was removed prior to collecting virus stocks. Virus stocks were quantified by plaque assay on Hela-MVR cells and sequenced by collecting infected 17Cl-1 or L929 cells using TRIzol™ Reagent (Invitrogen). The RNA from JHMV virions was likewise isolated from infected cells and submitted for full genome sequencing (SeqCenter).

### Virus Replication Assay

Cells were infected at the specified multiplicities of infection (MOIs) via a 1-hour adsorption phase with a minimum working volume of DMEM containing virus particles incubated at 37°C and periodically rocked to ensure complete coverage. Infected cells and supernatants were collected by freeze-thaw at −80°C. Viral titers were determined via plaque assay using HeLa-MVR cells.

### Plaque Size Measurements

JHMV was serially diluted and plated onto monolayers of indicated cell types in 24-well tissue culture plates. At 18hpi (DBT) or 20hpi (L929), cells were imaged using an Accu-Scope® EXI-310 with an attached Excelis™ HDS Digital Imaging System. Plaque diameters were measured using CaptaVision software calibrated with a stage micrometer.

### Plaque Reduction Assay

Stocks of JHMV *rep*D1329, D1329A, and N1347A were serially diluted and plated in triplicate on cell monolayers of HeLa, DBT, or L929 cells in 24-well tissue culture plates. Plaque formation was assayed at 20hpi.

### Viral Entry Assay

In a 24-well plate, DBT, L929, and HeLa cells were seeded at 2.25e5 cells/well and infected with a 100μl volume of MHV-JHM at an MOI of 1 using a 1-hour adsorption phase with regular shaking at 4°C. After the adsorption phase, cells were rinsed twice with PBS before fresh media was added to each well and plates were subsequently returned to the incubator at 37°C. At 2hpi, cells were removed from the incubator, rinsed with PBS, trypsinized, and the resulting cell suspension was transferred to 1.5ml microcentrifuge tubes. Tubes were centrifuged at 7,500 rpm for 5 minutes and the resulting cell pellets were resuspended in fresh PBS. Cells were rinsed in this fashion three times before a final resuspension in TRIzol™ Reagent (Invitrogen) frozen at −80°C. RNA extraction was carried out following the manufacturer’s recommended protocol.

### RT-qPCR Analysis

Cell monolayers were treated with ice cold TRIzol™ Reagent (Invitrogen) and RNA was extracted via the manufacturer’s recommended protocol. Synthesis of cDNA was conducted using random hexamers (Roche) with M-MuLV reverse transcriptase (NEB) via the manufacturer’s recommended instructions. Quantitation of cDNA transcripts was conducted using the Quantstudio 3 Real-Time PCR system (ThermoFisher) using the SYBR™ Green Universal Master Mix (ThermoFisher) with primers listed in **Table 2**.

**TABLE 2.**
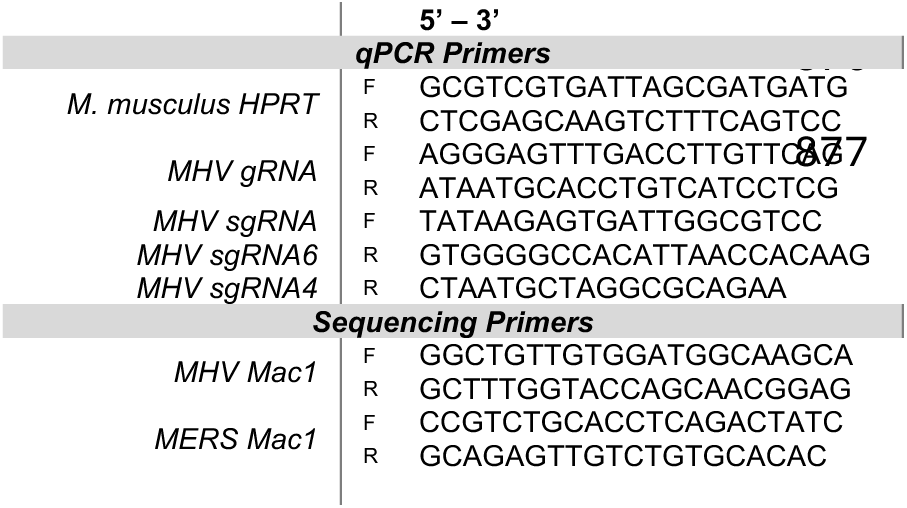
qPCR and Sequencing primers.

### Mouse Infections

For MHV mouse infections, 5-8 week-old male and female mice were anesthetized with isoflurane and inoculated intranasally with 1×10^4^ PFU recombinant JHMV or intraperitoneally with 500 PFU of A59. To obtain tissue for A59 virus titers mice were euthanized on different days post challenge, livers were removed and homogenized in phosphate buffered saline (PBS), and titers were determined by plaque assay on either Hela-MVR (MHV) cells.

### Immunoblotting

Total cell extracts were lysed in SDS-based lysis buffer containing Pierce™ Protease Inhibitor Cocktail (Thermo Scientific), PhosStop phosphatase inhibitor (Roche), 2% β-mercaptoethanol, and 1 mM PMSF. Proteins were resolved on a 10% SDS-PAGE gel, transferred to a polyvinylidene difluoride (PVDF) membrane, hybridized with a primary antibody, reacted with an infrared (IR) dye-conjugated secondary anti-body, visualized using a Li-COR Odyssey Imager (Li-COR), and analyzed using Image Studio software. Primary antibodies used for immunoblotting included anti-MHV N monoclonal antibody (1:10,000), anti-MHV S monoclonal antibody (1:5,000) (29), and anti-actin monoclonal antibody (1:10,000) (clone AC15; Abcam, Inc.). Secondary IR antibodies were purchased from Li-COR.

### Immunofluorescence Staining

Cells were seeded at approximately 1×10^5^ cells/ well in 8 Well Chamber, removable slides (ibidi) that were pre-coated with 50μg/ml collagen for 30 minutes. Cells were infected at an MOI of 0.5. At each time point, cells were rinsed with Hank’s Balanced Salt Solution with 0.01% Sucrose (HBSS/Su) and fixed with the addition of 2% paraformaldehyde in HBSS/Su. Permeabilization was conducted by rinsing 3x with HBSS/Su containing 0.1% Saponin (HBSS/Su/Sap) before blocking with 3% normal goat serum in HBSS/Su/Sap. Primary antibodies (mouse α-dsRNA clone rJ2, Millipore) (rabbit α-nsp3, in-house), each diluted 1:1,000 in HBSS/Su/Sap, were added at room temperature for 3 hours followed by rinsing 3× with HBSS/Su/Sap. Goat-derived secondary antibodies (AlexaFluor 488/555) diluted 1:200 in HBSS/Su/Sap were incubated 1 hour at room temperature protected from light followed by rinsing twice with HBSS/Su/Sap, once with HBSS/Su, and once with HBSS. Nuclear staining was achieved via the addition of 300nM DAPI in HBSS for 30 minutes followed by rinsing twice with HBSS. Slides were finalized with the addition of Vectashield® hardening Mounting Media (VectorLabs).

### Confocal Microscopy

Images were collected with a Leica DM6-SPE upright laser-scanning confocal microscope using a 63x oil-immersion objective lens with Type F oil. DAPI fluorescence was achieved with a 405 excitiation and 430/80 emission. Goat anti-mouse ALEXA 488 was imaged with 488 excitation 500/40 emission. Goat anti-rabbit ALEXA 555 was imaged with 561 excitation and 570/90 emission. Images were acquired using Leica LasX software.

### Quantification of Confocal Immunofluorescence

Quantification of protein staining was automated using a custom macro written in FIJI (ImageJ v.1.54f) (56). For each image, a median 3D filter was applied to multi-channel Z-stacks (x=1, y=1, z=1 for DBT; x=2, y=2, z=2 for BMDM) after which maximum intensity Z-projections were generated, and the channels were split. Regions of Interest (ROIs) were generated from these single-channel duplicates by automated thresholding via the Moments, Triangle, and Renyientropy methods for the DAPI, dsRNA/N protein, and nsp3 channels, respectively. Resulting binary masks were applied to the original Z-projections and fluorescence intensity of staining was measured within regions of interest (ROIs) on the relevant channel. The total area of each ROI was also recorded, and the Integrated Density (IntDen) displayed is a product of the mean fluorescence intensity and the ROI area.

### Colocalization Analysis

Manders’ coefficients were obtained using the BIOP JACoP plugin for FIJI (ImageJ v.1.54f) (56, 57). Median filters (x=1, y=1) were applied to maximum projection z-stacks, after which Triangle or RenyiEntropy auto thresholds were applied to the dsRNA and nsp3 channels, respectively. Resulting ROIs were cropped to produce proportions of total dsRNA signal overlapped (M1) and total nsp3 signal overlapped (M2).

### Protein Expression and Purification

Recombinant proteins were produced as previously described (35).

### *In vitro* de-MARylation Assay

MARylated PARP10 was generated via the incubation of purified GST-PARP10 with β-Nicotinamide Adenine Dinucleotide (β NAD^+^) (Millipore-Sigma) in a reaction buffer (50 mM HEPES, 150 mM NaCl, 0.2 mM DTT, and 0.02% NP-40). MARylated PARP10 was aliquoted and stored at −80°C. Purified SARS-CoV-2 Mac1 WT and D-A were co-incubated with MARylated PARP10 for indicated times at 37°C with a ratio of 1 μM Mac1 to 5 μM PARP10 in reaction buffer (50 mM HEPES, 150 mM NaCl, 0.2 mM DTT, and 0.02% NP-40). The reaction was halted with the addition of 2X Laemmli sample buffer containing 10% β-mercaptoethanol. Samples were then heated for five minutes at 95°C before loading into a Bolt™ 4%-12% Bis-Tris Plus Gel in MES running buffer (ThermoFisher). Direct protein detection was conducted via staining on a separate, identical gel using ReadyBlue® Protein gel stain (Sigma). Transfer to polyvinylidene difluoride (PVDF) membrane was achieved via the iBlot™ 2 Dry Blotting System (ThermoFisher). Immunoblotting of the MAR and GST signal was conducted with anti-mono ADP-ribose binding reagent MABE1076 (Sigma) and an anti-GST monoclonal antibody MA4-004 (ThermoFisher). Detection of primary antibodies was conducted using secondary infrared anti-rabbit and anti-mouse antibodies (LI-COR Biosciences). Samples were visualized using the Odyssey M® Imaging System (LI-COR Biosciences).

### Isothermal Titration Calorimetry

All ITC titrations were performed on a MicroCal PEAQ-ITC instrument (Malvern Pananalytical Inc., MA). All reactions were performed in 20 mM Tris pH 7.5, 150 mM NaCl using 100 μM of all macrodomain proteins at 25°C. Titration of 2 mM ADP-ribose or ATP (MilliporeSigma) contained in the stirring syringe included a single 0.4 μL injection, followed by 18 consecutive injections of 2 μL. Data analysis of thermograms was analyzed using one set of binding sites model of the MicroCal ITC software to obtain all fitting model parameters for the experiments. MERS-CoV and SARS-CoV-2 WT protein ITC data were previously published (35). These experiments were performed alongside the mutant proteins, thus serving as appropriate controls.

### Differential Scanning Fluorimetry

Thermal shift assay with DSF involved use of LightCycler® 480 Instrument (Roche Diagnostics). In total, a 15 μL mixture containing 8X SYPRO Orange (Invitrogen), and 10 μM macrodomain protein in buffer containing 20 mM Hepes, NaOH, pH 7.5 and various concentrations of ADP-ribose were mixed on ice in 384-well PCR plate (Roche). Fluorescent signals were measured from 25 to 95 °C in 0.2 °C/30-s steps (excitation, 470-505 nm; detection, 540-700 nm). Data evaluation and Tm determination involved use of the Roche LightCycler® 480 Protein Melting Analysis software, and data fitting calculations involved the use of single site binding curve analysis on GraphPad Prism.

### AlphaScreen Assay

The AlphaScreen reactions were carried out in 384-well plates (Alphaplate, PerkinElmer, Waltham, MA) in a total volume of 40 μL in buffer containing 25 mM HEPES (pH 7.4), 100 mM NaCl, 0.5 mM TCEP, 0.1% BSA, and 0.05% CHAPS. All reagents were prepared as 4X stocks and 10 μL volume of each reagent was added to a final volume of 40 μL. After 1h incubation at RT, streptavidin-coated donor beads (7.5 μg/mL) and nickel chelate acceptor beads (7.5 μg/mL); (PerkinElmer AlphaScreen Histidine Detection Kit) were added under low light conditions, and plates were shaken at 400 rpm for 60 min at RT protected from light. Plates were kept covered and protected from light at all steps and read on BioTek plate reader using an AlphaScreen 680 excitation/570 emission filter set. For data analysis, the percent inhibition was normalized to positive (DMSO + labeled peptide) and negative (DMSO + macrodomain + peptide, no ADPr) controls.

### Statistical Analysis

A Student’s *t* test was used to analyze differences in mean values between 2 groups, for multiple group comparisons, an ordinary one-way ANOVA was used. All results are expressed as means ± standard errors of the means (SEM) unless stated as standard differentiation (SD) or as geometric means ± 95% confidence internal. P values of ≤0.05 were considered statistically significant (*, P≤0.05; **, P≤0.01; ***, P≤0.001; ****, P ≤0.0001; ns, not significant).

### Data and Materials Availability

All data associated with this manuscript will be available through the data repository FigShare at 10.6084/m9.figshare.c.7755170 at the time of publication.

## ACKNOWLEDGEMENTS

We thank Ivan Ahel for providing protein expression plasmids, David Davido for critical reading, and the David Davido and Robin Orozco labs for helpful discussions. We thank our funding from the NIH, an NIH Graduate Training grant, the University of Kansas College of Liberal Arts and Sciences, and the Oklahoma Center for Advancement of Science & Technology.

## Author contributions

Conceptualization: JJOC, ARF

Data curation: JJOC, AR, RK, CK, YMA, ARF

Formal analysis: JJOC, AR, RK, CK, YMA, ARF

Funding acquisition: ARF

Methodology: JJOC, AR, RK, CK, NS, ARF

Investigation: JJOC, AR, RK, CK, YMA, XZ, XD, ARF

Project administration: ARF

Resources: PG, XD, ARF

Visualization: JJOC, AR, RK, CK, YMA, ARF

Validation: JJOC, AR, ARF

Supervision: XD, ARF Writing—original draft: JJOC, ARF

Writing—review & editing: JJOC, AR, RK, CK, NS, YMA, PG, XZ, XD, ARF

A.R.F. was named as an inventor on a patent filed by the University of Kansas for a live-attenuated SARS-CoV-2 vaccine

## SUPPLEMENTAL FIGURE LEGENDS

**Fig S1. MERS-CoV Mac1 D-A protein has poor ADP-ribose binding affinity. (A)** Isothermal titration caliorimetry of purified MERS-CoV WT D-A Mac1 protein incubated with ADP-ribose. *MERS-CoV WT protein data was previously published (27). **(B)** MERS-CoV Mac1 WT and DA proteins were tested for their ability to interact with an ADP-ribosylated peptide using an AlphaScreen assay. The results in A-B are from a single experiment representative of at least 2 independent experiments.

**Fig S2. JHMV-D1329A virus is defective in the production of viral genomic RNA (gRNA) and sub-genomic RNA6 (sgRNA6) compared to *rep*D1329 and N1347A viruses. (A-B)** L929 (A) and DBT (B) cells were infected with indicated viruses at an MOI of 0.1. Cells were collected at indicated times and genomic and RNAs were measured by qPCR using the *ΔΔ*CT method normalized to HPRT. Results are from one experiment representative of two independent experiments.

**Fig. S3. JHMV-D1329A produces nsp3 in the early stages of infection. (A)** DBTs were infected with indicated virus at an MOI of 0.5, fixed at 6 hpi, stained for nsp3, and analyzed by confocal microscopy. **(B)** Quantification of fluorescence signal displayed as a product of mean pixel intensity and total thresholded area (IntDen), representative of two independent experiments.

