## Supplemental Figures for "Mutations differentially affecting the coronavirus Mac1 ADP-ribose binding and hydrolysis activities indicate that it promotes multiple stages of the viral replication cycle"

**A**

| Macrodomain | Stoichiometry | Kd (μM) | ΔH (kcalmol$^{-1}$) | ΔG (kcalmol$^{-1}$) | -TΔS (kcalmol-1) |
|---|---|---|---|---|---|
| MERS-WT * | 1 | 7.2 ± 0.2 | -46 ± 0.7 | -29 ± 0.07 | 16 ± 0.6 |
| MERS DA | 1 | 279 ± 25 | -316 ± 27 | -25 ± 7 | 290 ± 33 |

**B**

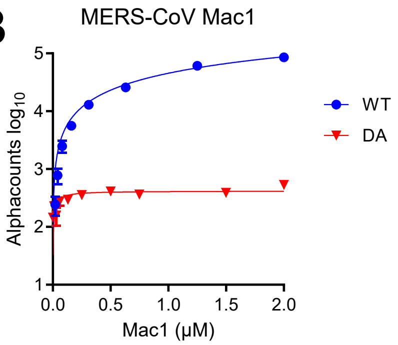


**Fig S1. MERS-CoV Mac1 D-A protein has poor ADP-ribose binding affinity. (A)** Isothermal titration caliorimetry of purified MERS-CoV WT D-A Mac1 protein incubated with ADP-ribose. *MERS-CoV WT protein data was previously published (27). **(B)** MERS-CoV Mac1 WT and DA proteins were tested for their ability to interact with an ADP-ribosylated peptide using an AlphaScreen assay. The results in A-B are from a single experiment representative of at least 2 independent experiments.

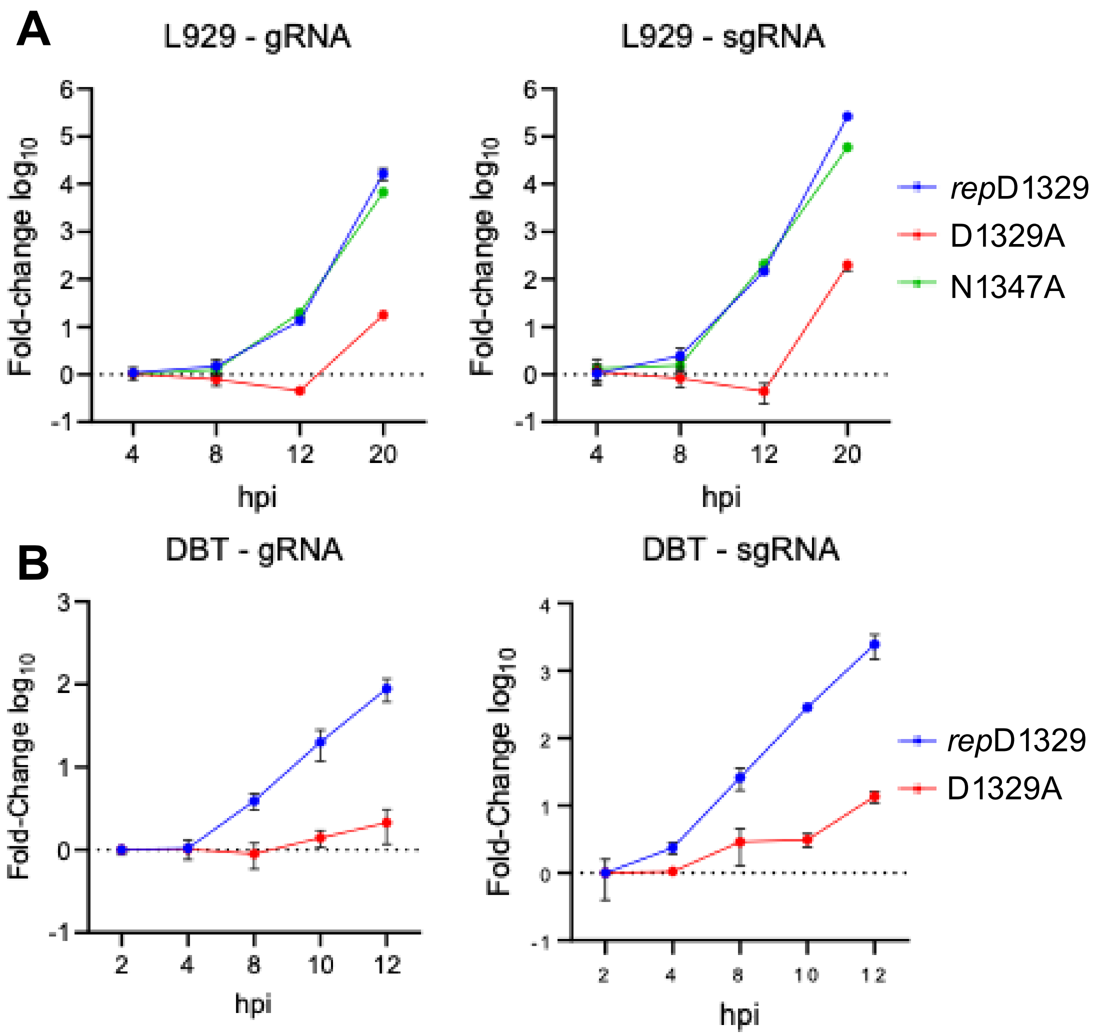


**Fig S2. JHMV-D1329A virus is defective in the production of viral genomic RNA (gRNA) and sub-genomic RNA6 (sgRNA6) compared to *rep*D1329 and N1347A viruses. (A-B)** L929 (A) and DBT (B) cells were infected with indicated viruses at an MOI of 0.1. Cells were collected at indicated times and genomic and RNAs were measured by qPCR using the *ΔΔ*CT method normalized to HPRT. Results are from one experiment representative of two independent experiments.

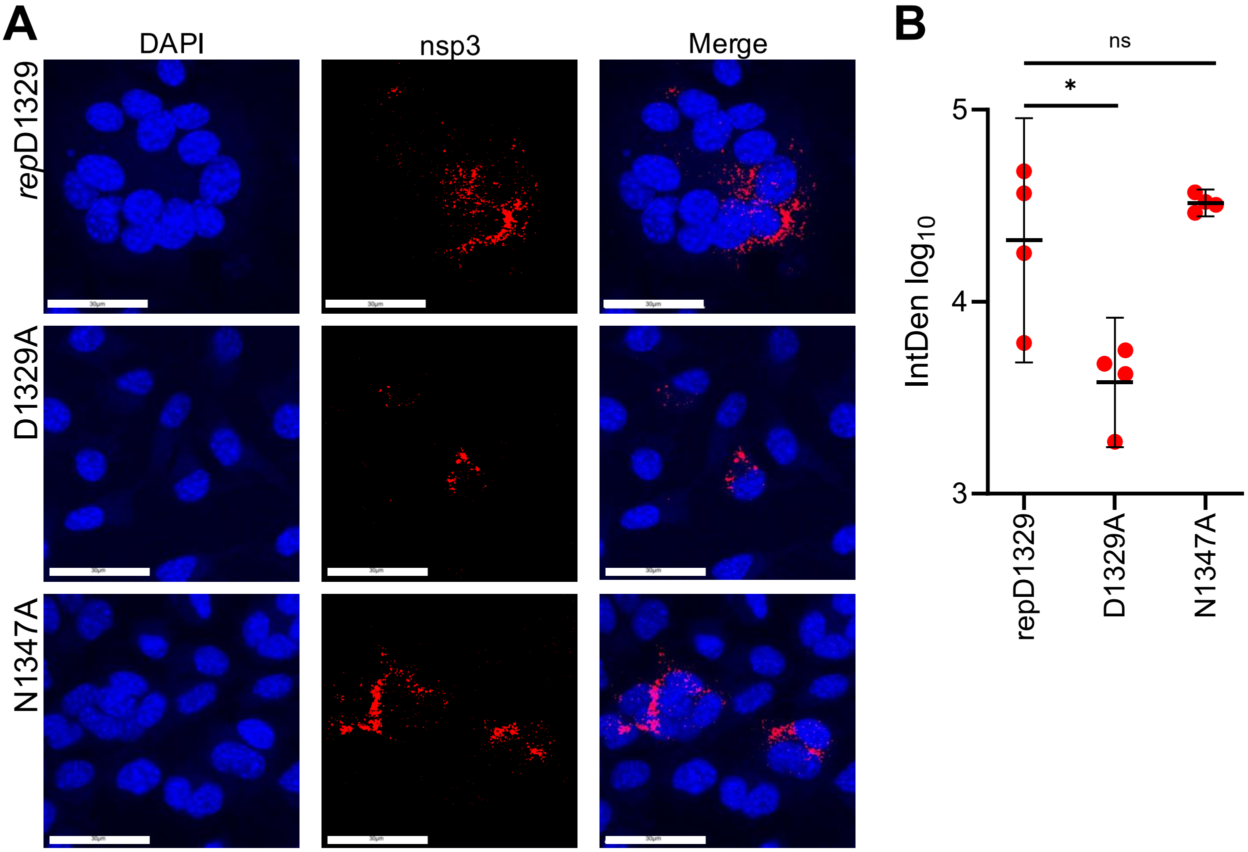


**Fig. S3. JHMV D1329A produces nsp3 in the early stages of infection. (A)** DBTs were infected with indicated virus at an MOI of 0.5, fixed at 6 hpi, stained for nsp3, and analyzed by confocal microscopy. **(B)** Quantification of fluorescence signal displayed as a product of mean pixel intensity and total thresholded area (IntDen), representative of two independent experiments.
